# DJ-1 glyoxalase activity makes a modest contribution to cellular defense against methylglyoxal damage in neurons

**DOI:** 10.1101/2022.02.18.481064

**Authors:** Melissa Conti Mazza, Sarah Shuck, Jiusheng Lin, Michael A. Moxley, John Termini, Mark R. Cookson, Mark A. Wilson

## Abstract

Human DJ-1 is a cytoprotective protein whose absence causes Parkinson’s disease and is also associated with other diseases. DJ-1 has an established role as a redox-regulated protein that defends against oxidative stress and mitochondrial dysfunction. Multiple studies have suggested that DJ-1 is also a protein/nucleic acid deglycase that plays a key role in the repair of glycation damage caused by methylglyoxal (MG), a reactive α-keto aldehyde formed by central metabolism. Contradictory reports suggest that DJ-1 is a glyoxalase but not a deglycase and does not play a major role in glycation defense. Resolving this issue is important for understanding how DJ-1 protects cells against insults that can cause disease. We find that DJ-1 reduces levels of reversible adducts of MG with guanine and cysteine in vitro. The steady-state kinetics of DJ-1 acting on reversible hemithioacetal substrates are fitted adequately with a computational kinetic model that requires only a DJ-1 glyoxalase activity, supporting the conclusion that deglycation is an apparent rather than a true activity of DJ-1. Sensitive and quantitative isotope-dilution mass spectrometry shows that DJ-1 modestly reduces the levels of some irreversible guanine and lysine glycation products in primary and cultured neuronal cell lines and whole mouse brain, consistent with a small but measurable effect on total neuronal glycation burden. However, DJ-1 does not improve cultured cell viability in exogenous MG. In total, our results suggest that DJ-1 is not a deglycase and has only a minor role in protecting neurons against methylglyoxal toxicity.

## Introduction

DJ-1 is a 20 kDa homodimeric protein that is conserved from bacteria to humans (1–3). In humans, mutations in DJ-1 (PARK7) cause rare forms of autosomal recessive parkinsonism (4). Eukaryotic DJ-1 promotes cell survival, particularly during oxidative stress or mitochondrial dysfunction, and is found in multiple cellular compartments (5–9). Consistent with its cytoprotective role, DJ-1 is highly expressed in several types of cancers (10–14) and it plays important roles in maintaining normal function in lung (15,16), eye (8,17), and kidney (18,19). A conserved cysteine (Cys106) in DJ-1 is both essential for its cytoprotective activity during oxidative stress (5,8,9,15,20,21) and is oxidation-prone, forming cysteine sulfinate (-SO_2_^-^) and sulfonate (-SO_3_^-^) species (5,22,23). The formation of Cys106-SO_2_^-^ has been proposed as one of several mechanisms that allow DJ-1 to act as a sensor of cellular redox state and to activate cytoprotective responses through the PTEN/Akt (24) and ASK1 (20,25) signaling pathways as well as by altering pathological protein aggregates (26–28). Other activities proposed for DJ-1, include protease (29,30), esterase (31), chaperone (32,33), transnitrosylase (34) and an RNA binding protein (35,36). Very recently, DJ-1 was shown to prevent protein damage by derivatives of 1,3, bisphosphoglycerate (37). Despite years of intensive study motivated by its biomedical importance, the molecular activities of DJ-1 that are responsible for its role in cell survival remain incompletely understood.

It has long been suspected that DJ-1 may possess an enzymatic activity. Cys106 has a low pK_a_ value of 5.4 and makes a hydrogen bond with a conserved protonated glutamic acid residue (Glu18) (38), which are features suggestive of an enzyme active site. These residues are conserved in the large DJ-1/PfpI superfamily (39), which contains well-established enzymes such as archaeal PfpI proteases (40,41), isocyanide hydratases (42–44), and the Hsp31 family of glutathione-independent glyoxalases (45). In all of these enzymes, the conserved cysteine residue is a catalytic nucleophile and the protonated glutamic/aspartic acid is a probable general acid/general base. In contrast to these validated enzymes, most proposed enzymatic functions of human DJ-1 have been controversial and confounded by weak apparent activities and variable assay conditions between studies (29,30,46,47).

Considerable interest has been generated by reports that human DJ-1 is a glutathione-independent glyoxalase that converts glyoxal to glycolate and methylglyoxal (MG) to L-lactate (48–51). In addition, DJ-1 has been reported to be a deglycase that repairs early glycation adducts that MG forms with DNA, RNA, small molecule thiols, and proteins (52–58). Both of these activities could protect cells by detoxifying reactive α-ketoaldehydes produced by metabolism, but in distinct ways. A glyoxalase removes MG directly, while a deglycase acts on glycated substrates and repairs primary damage, indirectly removing MG by repairing its initial adducts. The glyoxalase activity of DJ-1 has been widely reproduced but its reported k_cat_ is ~10^4^-10^5^ lower than the primary glutathione-dependent glyoxalase Glo1 (59,60). DJ-1’s deglycase activity has been controversial, and a recent study suggests that deglycation may be a secondary effect of DJ-1 acting on MG that is in rapid equilibrium with reversible MG adducts (61). Some reports indicate that DJ-1-mediated deglycation is important for repairing certain proteins (54,55,57,58,62), while others have indicated that DJ-1 has negligible effect on total cellular glycation burden (61,63) and does not deglycate disease-relevant proteins such as α-synuclein (26). In addition, several reports indicate that DJ-1 has no effect on cell viability under glycation stress in systems ranging from cultured human cells to yeast (51,61,63,64). Therefore, both the true substrate and physiological relevance of DJ-1’s anti-glycation action are debated (65).

Given the importance of DJ-1 in the etiology of several diseases, it is important to determine if a glyoxalase/deglycase activity is a major contributor to DJ-1 cytoprotection. If true, such an activity would have wide-reaching ramifications for the molecular etiology of several diseases. For example, a primary glyoxalase/deglycase activity for DJ-1 implies that reactive dicarbonyl species play a key role in the etiology of parkinsonism, as somatic DJ-1 deficiency invariably results in Parkinson’s disease in humans (4). A dominant glyoxalase/deglycase model for DJ-1 cytoprotection also implies that cells possessing enhanced ability to detoxify dicarbonyls are prone to neoplastic transformation, as DJ-1 is an oncogene and its levels are elevated in multiple cancers (14). Because even low levels of glycation can cause cellular dysfunction (66), quantitative methods for measuring macromolecular glycation such as isotope dilution mass spectrometry are needed to accurately evaluate the potential influence of DJ-1 on cellular glycation burden.

In this work, we use *in vitro* assays to show that DJ-1 cannot deglycate irreversible adducts of MG and guanine but can deglycate reversible adducts by an indirect mechanism. Steady-state enzyme kinetics using purified MG and reversible hemithioacetal substrates support a prior proposal that DJ-1’s apparent deglycase activity is due to its glyoxalase activity acting on free MG. Our kinetic constants are in reasonable agreement with those recently reported by Andreeva et al. (61) and confirm that DJ-1’s *in vitro* glyoxalase activity is many orders of magnitude lower than that of glyoxalase I (Glo1). Kinetic modeling demonstrates that the glyoxalase activity of DJ-1 is sufficient to explain the apparent deglycation kinetics. We use quantitative isotope-dilution mass spectrometry to show that DJ-1 measurably reduces the total DNA, RNA, and protein glycation burden in various neuronal cell lines and mouse brain, although the effect size is small. Cell survival in MG stress is unaffected by DJ-1 status, suggesting that DJ-1 plays a minor role in glycation defense.

## Results

### DJ-1 deglycates only reversible guanine-MG adducts

DJ-1 and its homologs have been reported to deglycate a range of MG adducts including lysine residues in proteins (55) and guanine bases in DNA and RNA (54) (Fig. 1A). While the role of DJ-1 on glycated amino acids have been studied by several groups with conflicting results (52,53,55,61,63,65), DJ-1’s influence on nucleotide glycation has been less studied (54). We investigated the role of DJ-1 in reversing MG-modified guanine nucleosides because this reaction has been proposed to play an essential role in maintaining genomic integrity in several organisms (54). MG glycation of deoxyguanosine (dG) in DNA can form either a reversible N^2^-(1,2-dihydroxy-2-methyl)ethano-2′-dG (cMG-dG) adduct or irreversible N2-(1-carboxyethyl)-2′-dG (CEdG) adduct via Schiff base formation and hydrolysis of an initial cMG-dG adduct (Fig. 1B). In situations where high concentrations of MG are present, CEdG can be further glycated to the stable MG-CEdG adduct (Fig. 1B). Glycation of RNA forms the corresponding guanosine adducts. The ability of recombinant DJ-1 to catalyze the deglycation of various guanine adducts was tested by mixing DJ-1 with a pre-equilibrated mixture of MG and dG and analyzing the resulting products by HPLC and mass spectrometry. We used MG purified by vacuum distillation (see Methods), as the purity of commercial MG is low (67). Addition of DJ-1 at the beginning of the reaction prevents formation of both reversible (cMG-dG) and irreversible (CEdG, MG-CEdG) adducts more effectively than when MG and dG are incubated together for one hour before DJ-1 addition (Fig. 2A; Supplemental Fig. S1). This reduction in MG adducts requires the catalytic nucleophile C106, as the C106S DJ-1 mutation abrogates this effect (Fig. 2B; Supplemental Fig. S1). In contrast to DJ-1’s ability to prevent formation of MG adducts when added early, incubation of purified R-CEdG and DJ-1 shows no change in R-CEdG chromatographic retention time on a C18 reverse phase column (67), indicating that this irreversible adduct is not a substrate for DJ-1 (Fig. 2C).

**Figure 1.**
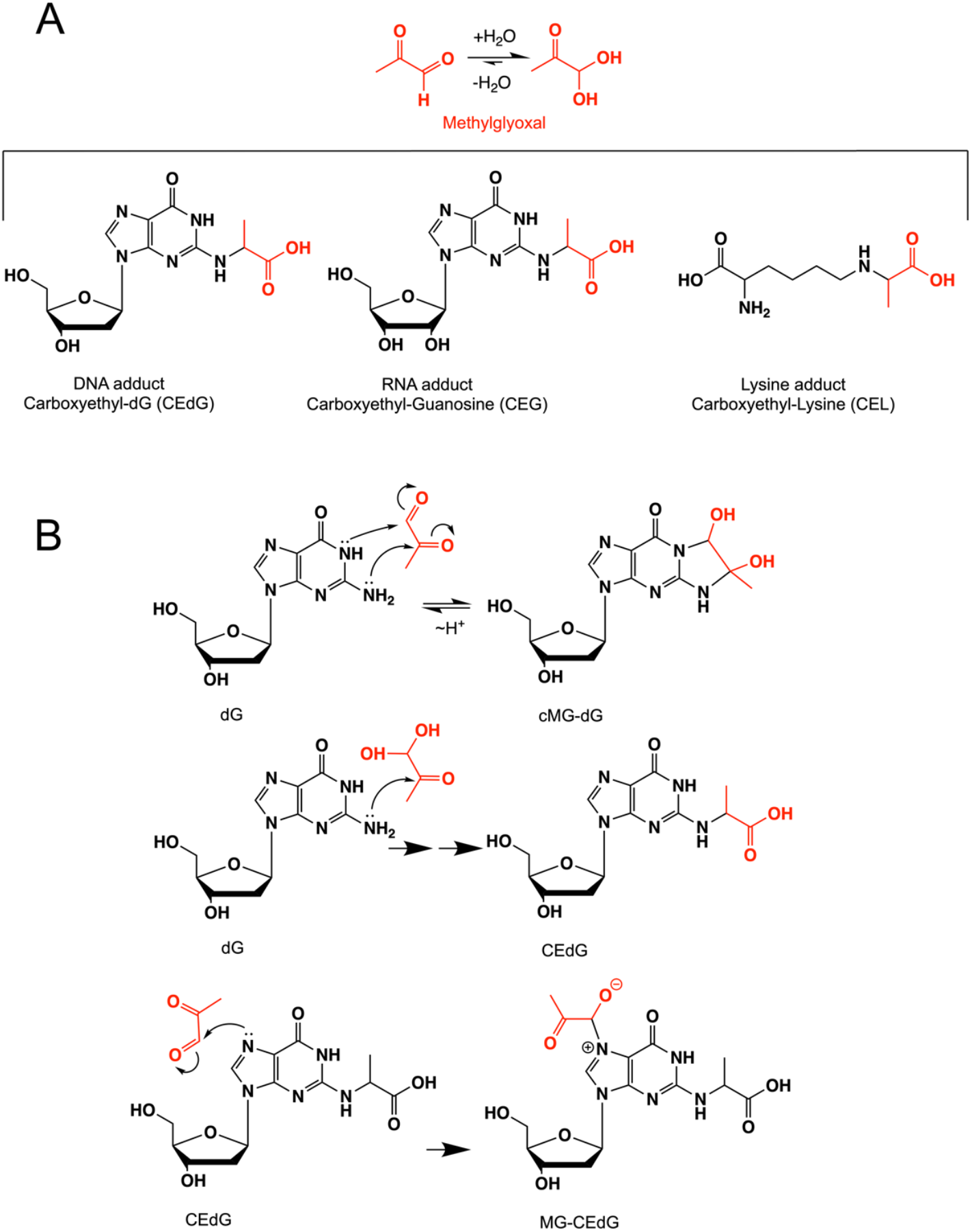
Adduct species generated by methylglyoxal (MG). (A) shows the hydration equilibrium reaction of MG (red), which lies to the right in aqueous solution. MG can form stable adducts with both guanine and lysine. (B) shows abbreviated mechanisms for MG modification of deoxyguanosine to form three adducts. Only cMG-dG is reversible, as indicated by arrows.

**Figure 2.**
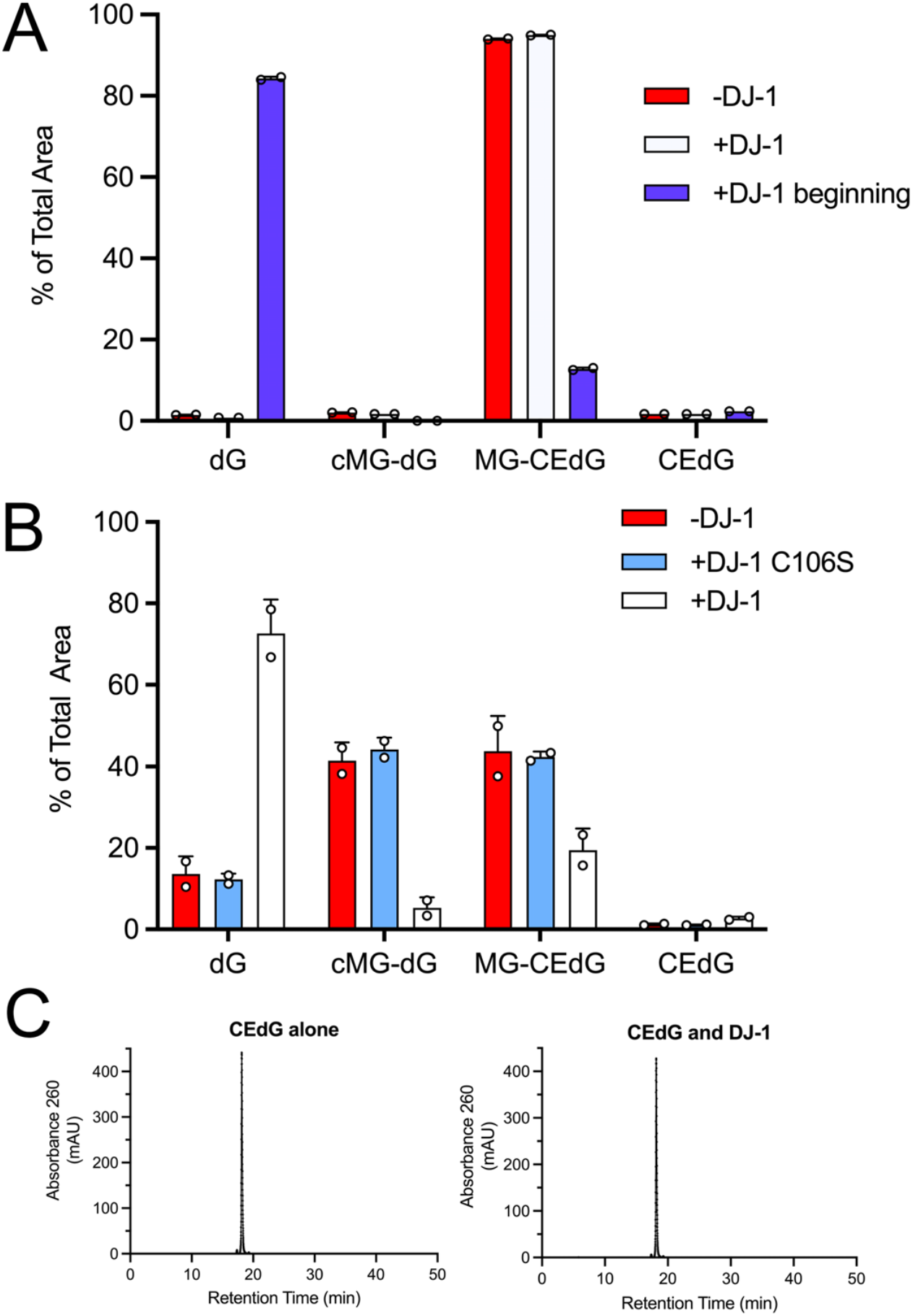
DJ-1 can only repair reversible adducts of MG and guanine in vitro. (A) Addition of DJ-1 at the same time as MG and dG (+DJ-1 beginning; blue) reduces levels of both reversible cMG-dG and irreversible MG-CEdG adducts, while addition of DJ-1 after one hour of preincubation of MG and dG (+DJ-1; white) results in an accumulation of irreversible MG-CEdG and cannot restore unmodified dG. (B) The C106S mutant (cyan), which removes the catalytic thiolate nucleophile, is inactive and cannot prevent dG modification. Only catalytically active DJ-1 prevents formation of dG adducts when added at the same time as MG. (C) Addition of DJ-1 results in no change in the HPLC retention time of purified CEdG, an irreversible adduct. In all panels, each measurement is shown as a circle with standard error of the mean shown in error bars.

By definition, reversible adducts of MG are in equilibrium with free MG. Our observations suggest that DJ-1 may act on free MG directly rather than on the early adducts of MG and guanine. If true, this predicts that a similar reduction of only the reversible MG adducts should be observed when small molecule aldehyde scavengers are added. We confirmed this by adding the aldehyde-reactive compounds DNPH and aminoguanidine, which behave similarly to DJ-1 in reducing the levels of reversible MG adducts with dG (Supplemental Fig. S2). In aggregate, these findings are consistent with DJ-1 acting on free MG as the primary substrate, which indirectly reduces the level of only the reversible MG-nucleoside adducts.

### DJ-1 has weak glyoxalase and apparent deglycase activities in vitro

Measurement of the putative deglycase activity of DJ-1 has produced variable and contradictory results, although the groups measuring this activity have used similar *in vitro* protocols (38,53,55,63). We measured the apparent deglycase activity of DJ-1 using a well-established assay that follows the decay of the 288 nm absorbance of the reversible hemithioacetal formed by incubation of MG with free thiols (68)(Fig. 3A). MG-hemithioacetals were formed by reaction of MG with N-acetyl-cysteine (NAC), glutathione (GSH), or coenzyme A (CoA) (see Methods). NAC is a standard model thiol, GSH is the most abundant small molecule thiol in the cell, and CoA has been used previously as a physiologically-relevant thiol whose glycation DJ-1 can reduce (56). Tris-containing buffers were not used to avoid previously reported buffer artifacts (63). Addition of DJ-1 reduces the concentrations of each of these three hemithioacetals with initial steady-state rates that are well-fitted using the Michaelis-Menten kinetics model (Fig. 3B,C). For the three hemithioacetal substrates, the apparent k_cat_ values were ~0.007-0.009 s^-1^ and K_M_ values were ~60-150 μM (Table 1). These activities are markedly lower than some previously reported values (55). DJ-1 apparent deglycase catalytic efficiency (k_cat_/K_M_) values are ~50-100 M^-1^ s^-1^, approximately 1×10^5^ fold lower than that of glyoxalase 1 acting on GSH-MG hemithioacetal as substrate (1.2 × 10^7^ M^-1^s^-1^ for E. coli GlxI) (69).

**Figure 3.**
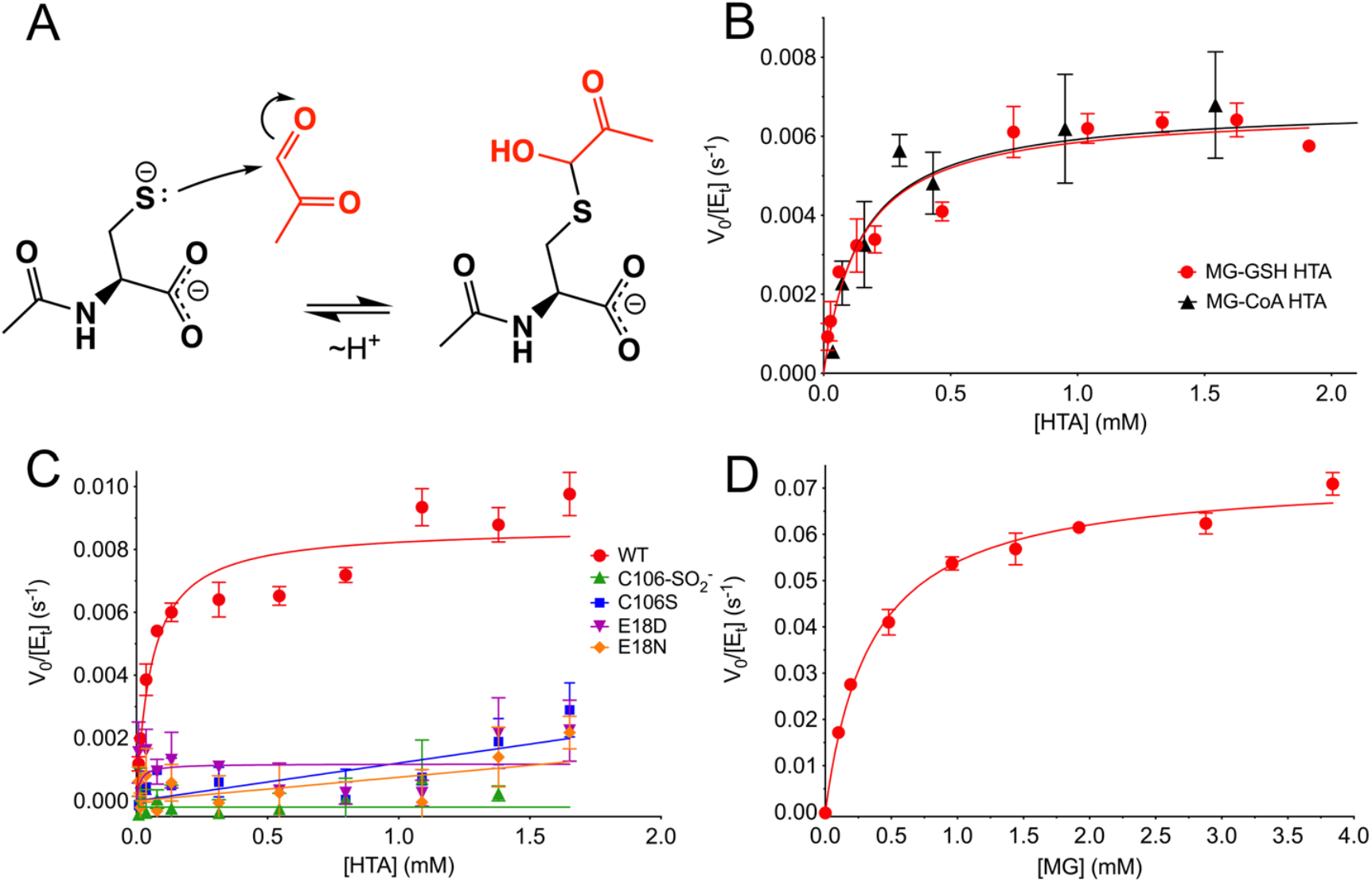
DJ-1 has apparent hemithioacetal deglycase and glyoxalase activities in vitro. (A) Mechanism of formation of hemiacetal formation by NAC (black) and MG (red). The hemithioacetal is reversible, as indicated by the arrows. (B) Steady-state enzyme kinetics of DJ-1 acting on the MG-glutathione (MG-GSH; red) and MG-coenzyme A (MG-CoA; black) hemitioacetal substrates. (C) Steady-state kinetics of DJ-1 acting on the MG-N-acetyl cysteine (MG-NAC) hemithioacetal substrate. Wild-type (WT) enzyme is in red, and mutants and oxidative modification are shown in the inset legend with the indicated symbols and colors. Only WT DJ-1 is active enough to be reliably fitted using the Michaelis-Menten model (solid lines). (D) Steady-state kinetics of DJ-1 glyoxalase activity against MG as substrate. In panels B-D, initial velocity (V0) is divided by total enzyme concentration ([Et]) on the Y-axis, rates were measured a minimum of three times with standard deviation shown in bars, and the fitted Michaelis-Menten curves are shown as solid lines.

**Table 1.**
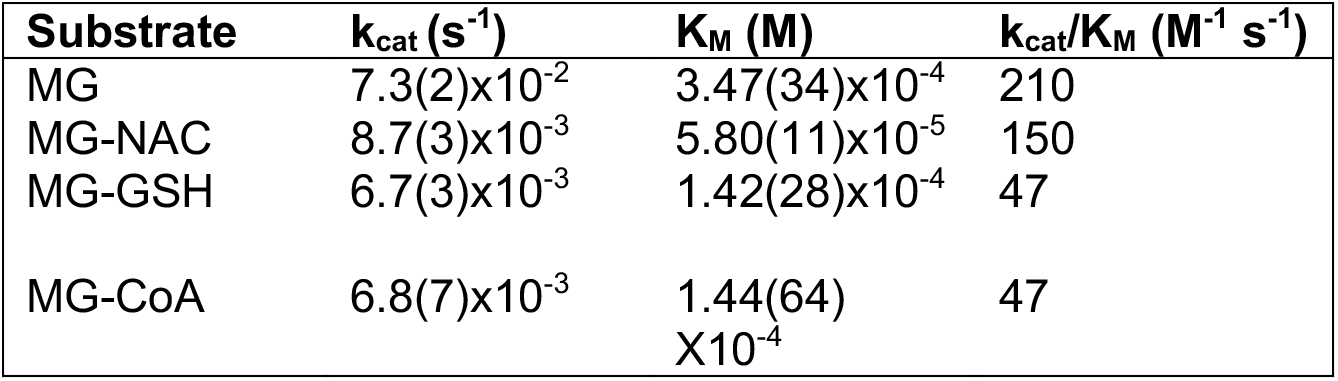
DJ-1 kinetic parameters from experiment.

The reactive Cys106 residue is the presumed catalytic nucleophile in human DJ-1. We confirmed that DJ-1’s ability to consume MG-NAC hemithioacetal is Cys106-dependent by showing that neither the C106S mutant nor the Cys106-sulfinate (C106-SO_2_^-^) oxidized form of the protein has significant apparent deglycase activity (Fig. 3C). The active site environment of Cys106 features a protonated Glu18 residue that makes a hydrogen bond to the Cys106 thiolate (38) and was identified as catalytically important in prior reports of DJ-1 glyoxalase activity (48). Consistent with these prior reports, mutations at the nearby protonated Glu18 ablated DJ-1-mediated consumption of MG-NAC hemithioacetal (Fig. 3C). Our prior structural studies showed that E18D, E18N, and wild-type DJ-1 have nearly identical crystal structures and that Glu18 mutations only slightly perturb the steric environment of Cys106 (38,70), suggesting that these mutations do not eliminate DJ-1’s apparent deglycation activity *via* major changes in protein structure. Instead, it is likely that mutations at Glu18 remove an important general acid/base in the proposed DJ-1 glyoxalase mechanism (Supplemental Fig. S3).

Because hemithioacetals are obligatorily reversible MG adducts that cannot be converted to irreversible glycated products via the Amadori rearrangement, they always exist in equilibrium with free MG. Therefore, DJ-1’s apparent deglycation activity may be due to its action on free MG, as we proposed above for guanine MG adducts and was also proposed by others for other reversible MG adducts (61). DJ-1 has an established glyoxalase activity, although estimates for its kinetic parameters have varied markedly (48,50,55,61,63,71). Unlike some other glyoxalases in the DJ-1 superfamily, human DJ-1 produces exclusively L-lactate as a product, permitting measurement of its full catalytic activity using an L-lactate-coupled assay (50,71). We measured the rate at which DJ-1 converts MG to L-lactate in real time using L-lactate oxidase, horseradish peroxidase, and resorufrin in a coupled assay that is similar to one used previously (61)(Methods). Varying the concentration of the coupling enzymes L-lactate oxidase and horseradish peroxidase has little effect on the initial rate measurements, confirming that DJ-1’s activity is rate-limiting in this coupled assay. DJ-1’s glyoxalase activity is well-fitted using the Michaelis-Menten model (Fig. 3D) and produces kinetic constants (Table 1) which are in reasonable agreement with the k_cat_~0.02 s^-1^ reported by Andreeva et al. (61). DJ-1’s glyoxalase k_cat_ is ~10-fold greater than its apparent deglycase k_cat_.

### Kinetic modelling indicates that DJ-1 glyoxalase activity is sufficient to explain its apparent deglycase activity

To determine if the low apparent deglycase activity for DJ-1 could be explained by its higher glyoxalase activity, we used kinetic modeling. We simulated the DJ-1-catalyzed glyoxalase reaction using a simple Michaelis-Menten mechanism (reaction 1) coupled to hemiothioacetal production via the reaction of methylglyoxal with N-acetylcysteine (Scheme 1).

**Scheme 1.**
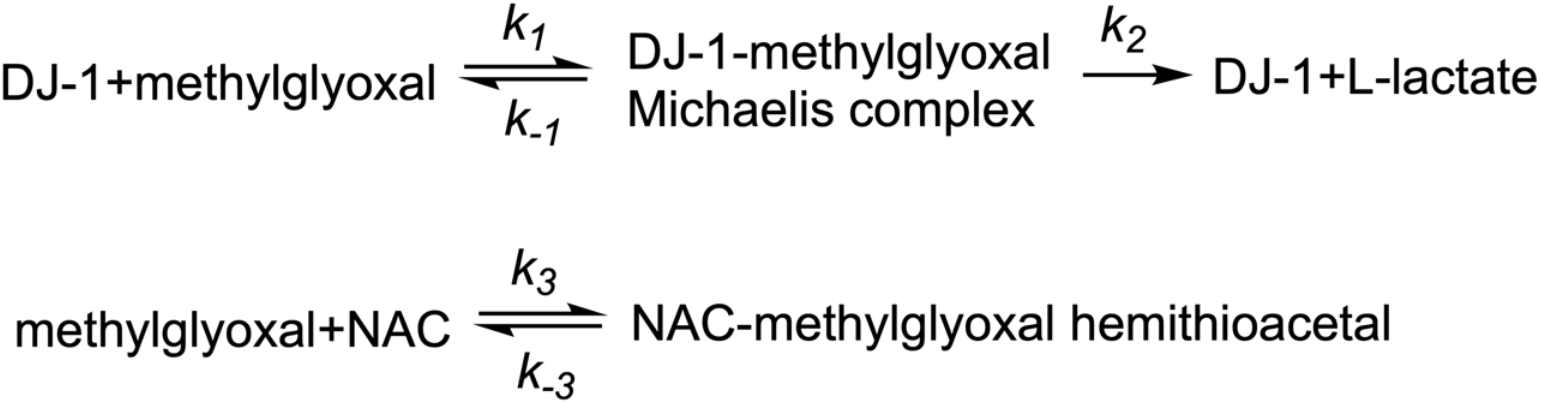

The measured progress curve for DJ-1-mediated loss of NAC-MG hemithioacetal can be adequately modeled using a kinetic scheme that includes only a glyoxalase activity for DJ-1 and an equilibrium of free MG and MG-NAC hemithioacetal, without need for a DJ-1 deglycase activity (Fig. 4A). The modeled kinetic parameters for DJ-1 glyoxalase activity also agree well with the measured glyoxalase activity progress curves of the enzyme (Fig. 4B) and produce glyoxalase k_cat_ and K_m_ values that agree well with the experimentally determined ones (Table 2). The ability to fit the data in Fig. 4 without needing to invoke a postulated DJ-1 deglycase activity supports the conclusion (61) that DJ-1 is not a primary deglycase but reduces the concentrations of reversible MG adducts through its action on MG and the equilibria in Scheme 1.

**Figure 4.**
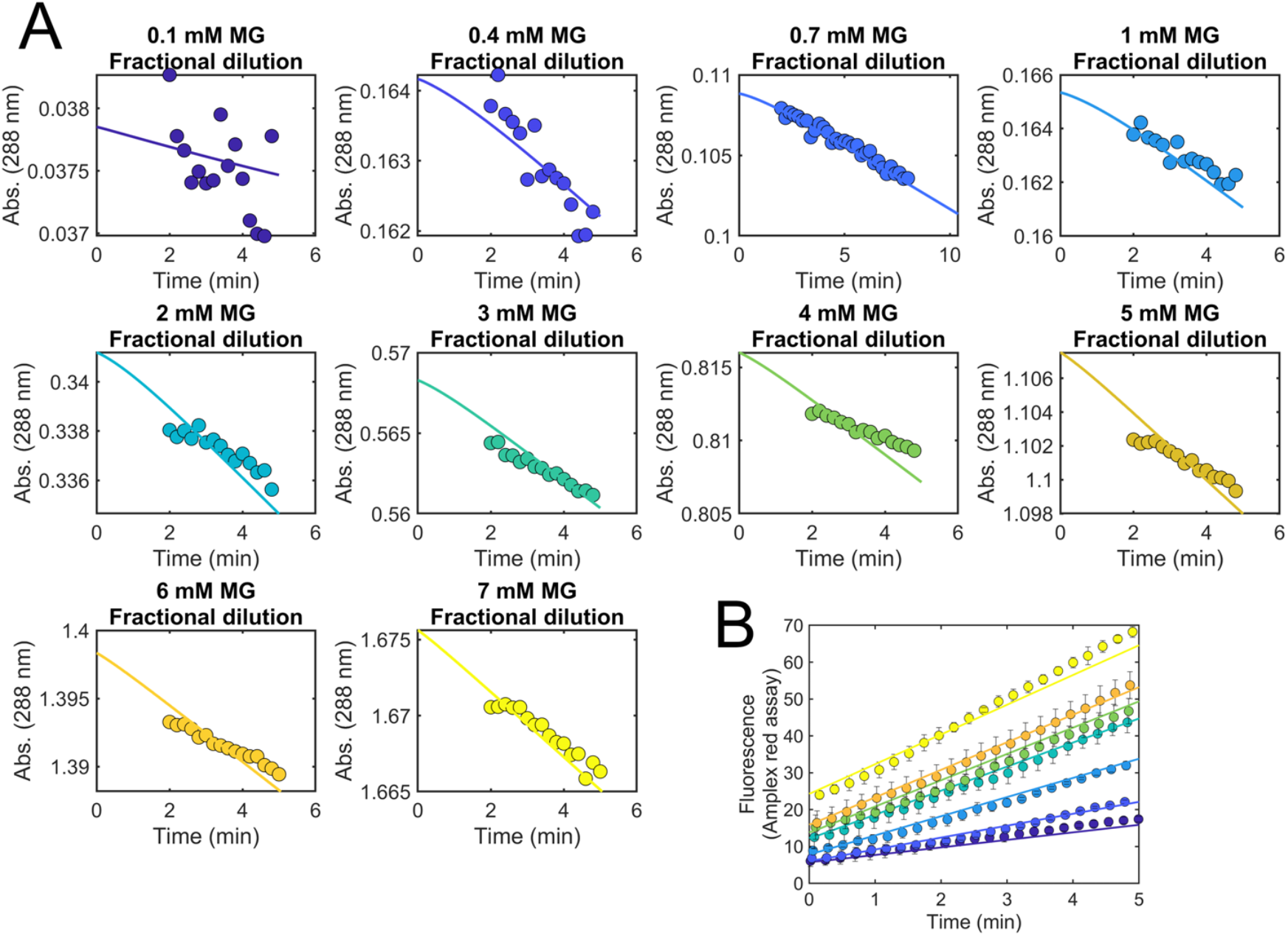
Kinetic simulation of DJ-1 deglycase and glyoxalase activities. (A) Experimental progress curves for loss of MG-NAC hemithioacetal signal at 288 nm (colored dots) are superimposed with kinetic modeling results (solid lines) for various concentrations of total MG concentration at initial dilution. The simulation requires only a DJ-1 glyoxalase activity to produce good fits to the apparent deglycase activity. (B) Experimentally measured DJ-1 glyoxalase activity against various concentrations of MG (colored dots) is well-modeled by the kinetic simulation (solid lines) whose rate constants also explain the apparent deglycase activities in (A).

**Table 2.** DJ-1 kinetic parameters with MG substrate from kinetic simulation.

<sup>a</sup>Lower and upper boundaries were determined by a published algorithm that varies the rate constants to find values that fall within a 0.83 relative error to the
| Rate constant | Best-fit value | <sup>a</sup>Lower boundary | <sup>a</sup>Upper boundary |
| --- | --- | --- | --- |
| $k_1$ (M<sup>-1</sup> sec<sup>-1</sup>) | 360 | 123.83 | 11,466.67 |
| $k_{-1}$ (sec<sup>-1</sup>) | 7.82x10<sup>-2</sup> | 2.62 x 10<sup>-12</sup> | 4.13 |
| $k_2$ ($k_{cat}$) (sec<sup>-1</sup>) | 4.02x10<sup>-2</sup> | 3.82x10<sup>-2</sup> | 4.27x10<sup>-2</sup> |
| <sup>c</sup>Calculated $K_M$ (M) | 3.29x10<sup>-4</sup> | <sup>d</sup>2.76x10<sup>-4</sup> | <sup>d</sup>3.63x10<sup>-4</sup> |
| $k_3$ (M<sup>-1</sup> sec<sup>-1</sup>) | 3.25 | 1.98 | 104.17 |
| $k_{-3}$ (sec<sup>-1</sup>) | 6.5x10<sup>-3</sup> | <sup>b</sup>ND | <sup>b</sup>ND |

best-fit (82).
<sup>b</sup>KinTek global explorer software only estimates one rate constant when the ratio is being constrained by the equilibrium constant (500 M<sup>-1</sup>).
<sup>c</sup>The $K_M$ was calculated using the definition $\left( K_m = \frac{k_{-1} + k_2}{k_1} \right)$
<sup>d</sup>Lower and upper boundaries are not determined directly in KinTek global software for steady-state constants therefore they were calculated from the collection of rate constants that fit the data within a 0.83 relative error from the best-fit.

### DJ-1 reduces total protein and nucleic acid glycation burden in cultured neuronal cells and mice

DJ-1’s contribution to reducing the overall glycation burden in cells is contentious, with some studies indicating major effects (54,55,58) and others detecting no changes (61,63). In these studies, protein glycation has been measured primarily using Western blotting. Western-based detection of glycated macromolecules is complicated by the need for specific antibodies against the modification of interest and the difficulty of quantifying the blots. To address these limitations, we used quantitative isotope dilution mass spectrometry with internal ^15^N_5_-CEdG,^15^N_5_-CEG, CEL-d4 and Lys-d4 standards for quantifying glycated guanine in DNA, RNA and glycated lysine in proteins. Using dopaminergic M17 neuroblastoma cells, we knocked down DJ-1 using siRNA (see Methods) and observed a trend of increased CEdG, CEG, and CEL levels when DJ-1 levels are decreased, although these effects are small (Fig. 5A).

**Figure 5.**
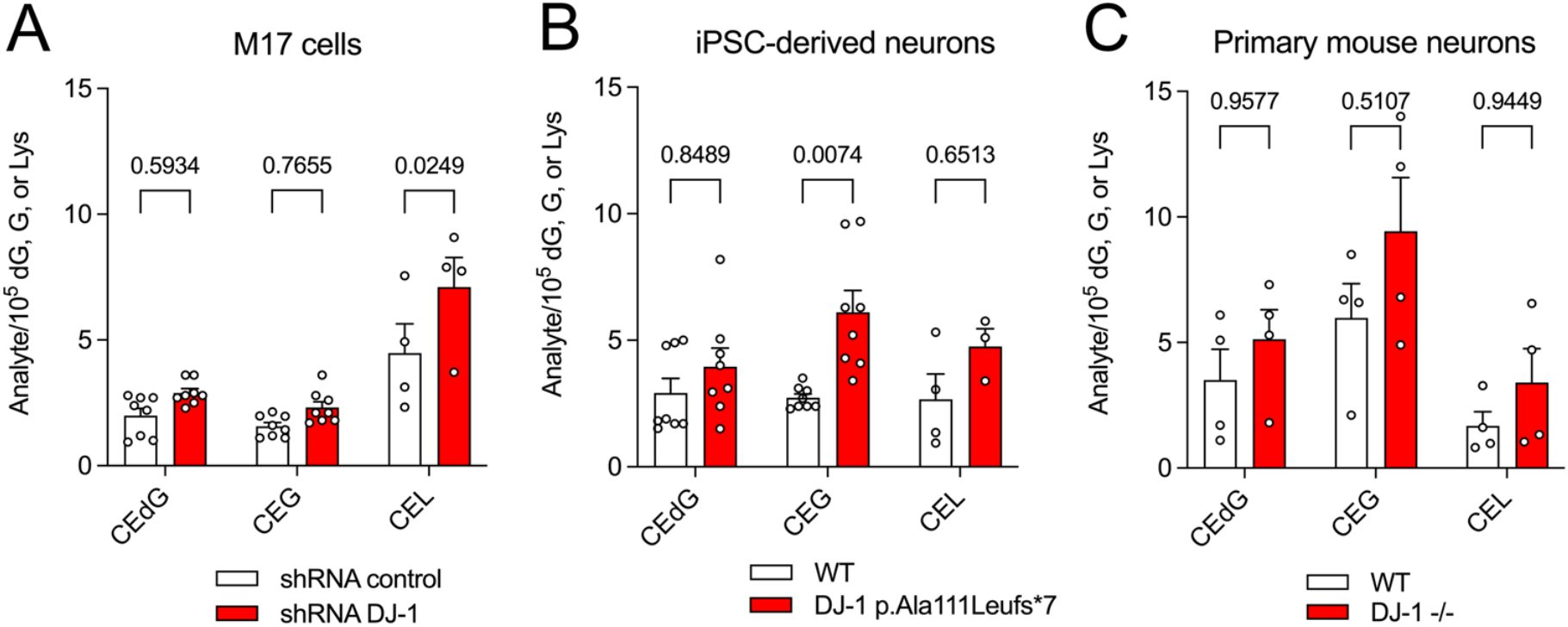
DJ-1 has a modest impact on cellular concentrations of irreversible glycation products in neurons. In all panels, isotope-dilution mass spectrometry was used to obtain relative concentrations of modified vs. unmodified dG, G, or Lys. Two-way ANOVA was used for statistical analysis with p values shown. Small but consistent elevations in glycated products were observed in (A) siRNA knockdown of DJ-1 in immortalized M17 neuroblastoma, (B) iPSC-derived neurons derived from wild-type (WT) cells and cell bearing a DJ-1 missense mutation that eliminates protein (DJ-1 p.Ala111Leufs*7) and (C) primary neurons from WT and DJ-1^-/-^ mice. Each measurement is shown as a circle with standard error of the mean shown in error bars.

Although M17 cells are biochemically similar to the vulnerable neurons in Parkinson’s Disease, they are immortalized and thus have potential alterations in metabolism that could affect glycation. To address this, we also measured glycated products in two distinct neuronal lineages: induced pluripotent stem cell-derived forebrain neurons created from fibroblasts donated by a patient bearing a A111L missense mutation that eliminates steady-state DJ-1 (72) and primary neurons cultured from DJ-1^-/-^ mice. In both DJ-1-deficient neuronal cultures, glycation products are slightly elevated (Fig. 5A-C), although this increase only reaches the p<0.05 significance threshold for CEL in M17 cells and CEG in iPSC-derived neurons (Fig. 5A,B). A similar effect is seen in overall mouse brain tissue, where DJ-1 knockout elevates all three classes of glycated products by 1-2 modified species/10^6^ unmodified, which is statistically significant for CEG and CEL (Supplemental Fig. S4).

The glutathione-dependent glyoxalase I/II (GloI/II) system is the primary means for detoxifying MG in the cell. Because the *in vitro* catalytic rate of GloI is several orders of magnitude greater than that of DJ-1, it is possible that DJ-1 would make a larger contribution to glycation defense when the GloI/II is impaired. We tested this using buthionine sulfoximine (BSO), an inhibitor of γ-glutamyl cysteine ligase, the rate-limiting enzyme in glutathione (GSH) biosynthesis (73). As GSH is a co-substrate for GloI, reduction in the GSH pool should reduce GloI activity in cells. BSO administration to M17 neuroblastoma cells markedly increases CEL levels when compared to vehicle alone, although the effect on CEG and CEdG levels are not significant. When BSO is applied to DJ-1 knockdown cells, it leads to significant increases in all three glycated metabolites (CEdG, CEG, and CEL) and this effect is again most pronounced for CEL (Fig. 6C). The enhancement of glycation when BSO is applied to DJ-1 knockdown cells suggests that Glo1’s stronger glyoxalase activity dominates the cellular MG defense compared to DJ-1’s weaker glyoxalase activity.

**Figure 6.**
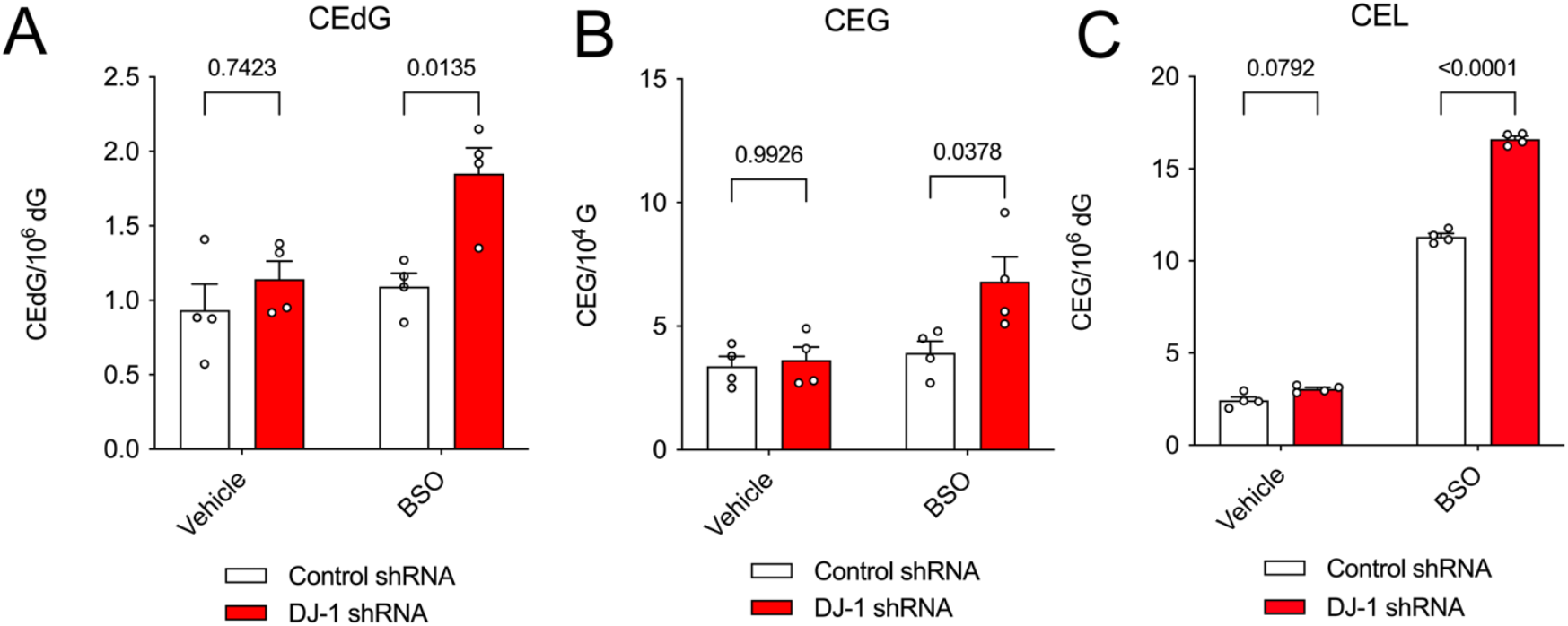
Chemical reduction in cellular glutathione enhances the glycation burden of DJ-1 knockdown M17 cells. In all panels, isotope-dilution mass spectrometry was used to obtain relative concentrations of modified vs. unmodified dG, G, or Lys in cultured M17 neuroblastoma cells. Two-way ANOVA was used for statistical analysis with p-values shown. BSO was used to reduce cellular levels of GSH, which is a co-substrate for the dominant glyoxalase Glo1. Administration of BSO enhances the effect of DJ-1 deficiency on CEdG (A), CEG (B) and CEL (C) levels. Each measurement is shown as a circle with standard error of the mean shown in error bars.

### DJ-1 does not protect cells against MG toxicity

DJ-1 decreases total cellular CEdG, CEG, and CEL levels in some instances, although the absolute magnitude of this decrement is relatively small (~1-2 glycated products/10^5^ unmodified). To determine if this reduction in glycation corresponds to increased cellular protection against MG toxicity by DJ-1, we measured the effect of DJ-1 on survival of both M17 neuroblastoma cells and iPSC-derived neurons in 10-10000 μM exogenously added MG. M17 cells show a clear loss of viability with an IC_50_ of 809.2 μM MG, but DJ-1 siRNA knockdown has no effect on cell survival (Fig. 7). Sensitization to MG, resulting in a decrease of IC_50_, would be expected if DJ-1 made a significant contribution to cellular viability during MG challenge of these cells. Neurons derived from iPSCs show no loss of viability across a 10-1000 μM MG range, and the inactivating A111L DJ-1 mutation does not sensitize them to MG (Fig. 7). These negative results are consistent with prior reports that loss of DJ-1 has no effect on MG sensitivity in Drosophila and HEK293 cells (61,63).

**Figure 7.**
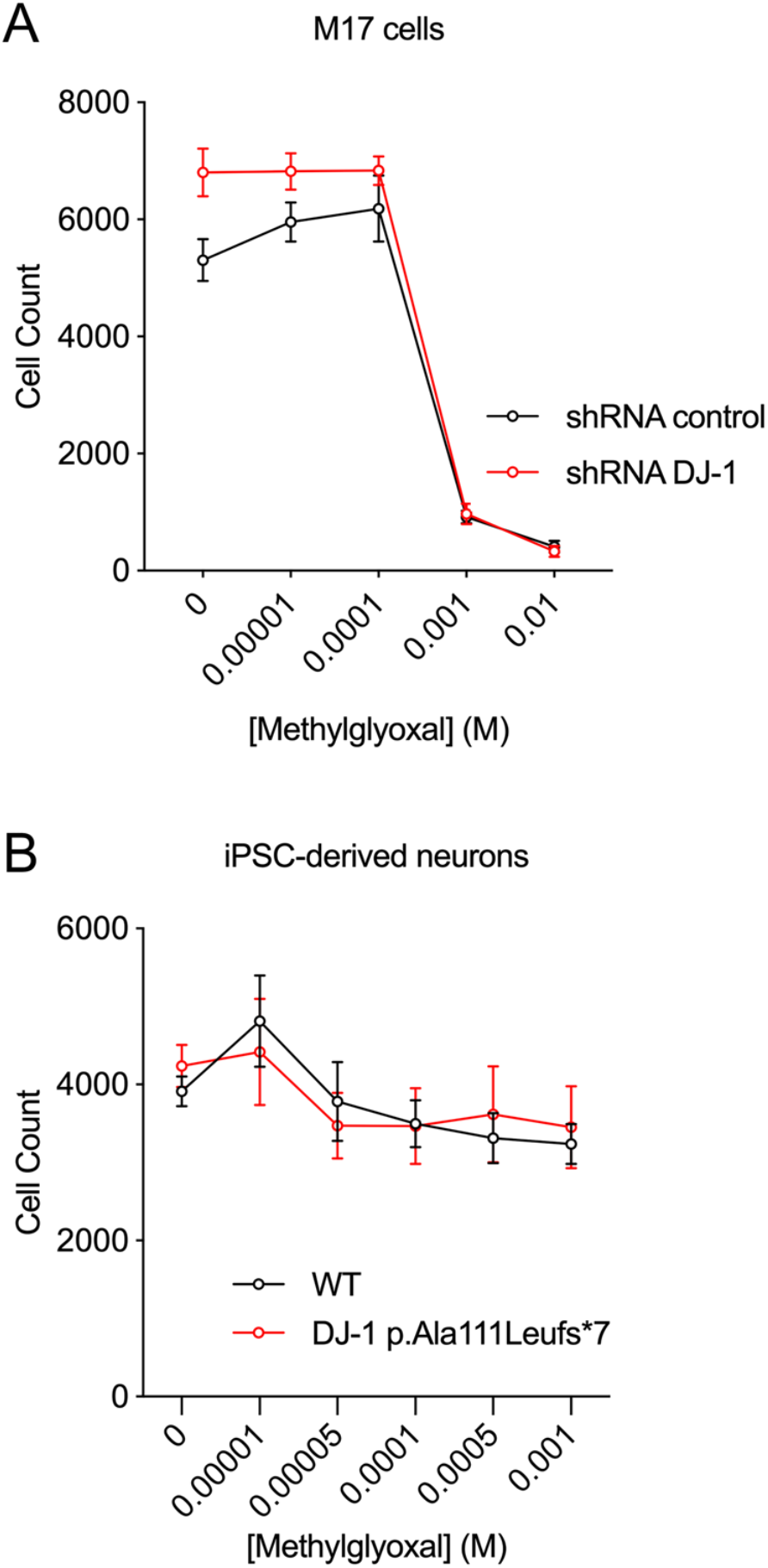
DJ-1 deficiency does not enhance cell death in the presence of exogenous MG. Exogenous MG was added to cultured M17 cells (A) and iPSC-derived neurons (B). No enhancement of cell death was observed when DJ-1 was knocked down with siRNA (A) or absent at the protein level (B).

## Discussion

DJ-1 defends multiple types of cells against oxidative stress and mitochondrial damage. One model of DJ-1-mediated cytoprotection is that it is a redox-responsive protein that uses cysteine oxidation to sense changes in redox homeostasis and enhances cell survival through the PTEN/Akt (74), Nrf2 (16,75), and ASK1 (20,25) signaling pathways. However, DJ-1 has more recently been reported to have Cys106-dependent glyoxalase and deglycase activities, indicating that the protein may also play important roles in defense against reactive carbonyl species such as MG. DJ-1’s glyoxalase activity is widely corroborated, although it is significantly less efficient than Glo1 and there is considerable disagreement regarding DJ-1’s kinetic constants. The deglycase activity for DJ-1 is more controversial, with some groups reporting that it is either an artifact (63) or an apparent activity of DJ-1 acting on methylglyoxal (61), while others report that deglycation is the most important activity of DJ-1 (54,55).

Our results help resolve this dispute, as we find that DJ-1 is a weak glyoxalase with a k_cat_~0.07 s^-1^. This is ~100x lower than the average enzyme k_cat_ value (76), but is still 10x higher than the maximal rate of DJ-1’s apparent deglycase activity. There has been disagreement in the literature about kinetic constants for DJ-1’s glyoxalase activity (65), and we note that our results agree reasonably well with those of Andreeva et al. measured in the same buffer (61). We observe that DJ-1 can only reduce the levels of freely reversible MG adducts in vitro, and our kinetic modeling shows that DJ-1’s glyoxalase activity is sufficient to explain its this apparent activity on MG adducts. Here again, these new quantitative kinetic modeling results support the conclusions of Andreeva at al. (61) that the apparent deglycase activity of DJ-1 is likely due to its action on free MG regardless of the reversibly glycated species: aminocarbinol, hemithioacetal, cyclic dihydroimidazolone, or others. This conclusion contradicts several reports from the Richarme group (52–55) but is chemically sensible, as there is no clear mechanism by which a cysteine nucleophile-dependent enzyme could deglycate these diverse species, all of which lack a plausible electrophilic center for Cys106 thiolate attack. In addition, we find that both the E18N and E18D mutants eliminate DJ-1’s ability to revert reversibly glycated species, consistent with some prior reports (48,56) and demonstrating that Glu18 is catalytically essential for DJ-1 glyoxalase activity. Considering these results in the context of the DJ-1 crystal structure, we propose a mechanism where the protonated Glu18 is the general acid that protonates the oxyanion formed upon Cys106 attack at the electrophilic aldehyde carbon atom of MG (Supplemental Fig. S3).

Prior studies show that E18N and E18Q DJ-1 can protect cells from mitochondrial damage induced by rotenone, preserve normal mitochondrial morphology (70), and are functional in other assays of DJ-1 activity (21,25). These results indicate that aspects of DJ-1 cytoprotection related to oxidative stress response and mitochondrial dysfunction are not explained by DJ-1’s glyoxalase activity. Furthermore, any cytoprotective enzymatic activity that requires Cys106 as the catalytic nucleophile will be abrogated by oxidation of Cys106 that occurs during oxidative stress, suggesting that DJ-1’s known enzymatic and redox-sensing protective mechanisms are orthogonal and require different cellular pools of the protein.

The low catalytic efficiency of DJ-1’s glyoxalase activity (k_cat_/K_M_=210 M^-1^s^-1^) raises questions about its physiological relevance, as it is much lower than the ~10^7^ M^-1^ s^-1^ k_cat_/K_M_ of Glo1 (69,77). Glo1, a highly conserved and widely expressed enzyme that converts the MG-glutathione hemithioacetal to S-lactoylglutathione, is the dominant mechanism of MG detoxification in most cells. It seems implausible that DJ-1 could contribute significantly to the cellular defense against MG against a background of more catalytically proficient Glo1 unless Glo1 activity is diminished by glutathione depletion or other mechanisms. Despite these considerations, we find using sensitive and quantitative isotope dilution mass spectrometry that the absence of DJ-1 results in small elevations in neuronal CEdG, CEG, and CEL, three surrogate biomarkers for MG reaction with DNA, RNA, and protein, respectively. These elevated levels are found in immortalized M17 neuroblastoma cells, iPSC-derived neurons, primary mouse neurons, and whole mouse brain, establishing that this is observed across multiple neuronal types. Moreover, the effect is observed both in vivo and under standard cell culture conditions without external (and unphysiological) administration of bolus MG. Although the effect size is modest, it is nonetheless surprising given the low catalytic activity of DJ-1. We offer two speculative hypotheses that may explain this result. First, DJ-1 may have substantially greater glyoxalase activity in the cell than observed in vitro using recombinant protein, perhaps owing to the presence of yet-unknown modifications or regulators in the cell that enhance its activity. This explanation may seem contrived, however this effect has been observed for the related Hsp31 glyoxalase in *Saccharomyces cerevisiae*, although the cause of endogenous Hsp31’s enhanced activity is unknown (78). Further supporting the sensitivity of DJ-1’s glyoxalase activity to the details of the solution reaction environment, the in vitro DJ-1 glyoxalase activity is markedly higher in phosphate buffer than in PBS (79). Second, it is possible that DJ-1’s glyoxalase activity is not directly responsible for its ability to reduce cellular glycation burden. In this model, DJ-1 would enhance the activities of other pathways that are more effective at reducing steady-state levels of MG. This model could be tested using mutations such as E18N that eliminate DJ-1 glyoxalase activity but preserve its protective activity against oxidative and mitochondrial damage stressors and warrant further investigation.

DJ-1 does not improve cellular viability against MG toxicity in our experiments, supporting some prior reports (61,63) while contradicting others (48,55). It is possible that DJ-1’s modest ability to reduce glycation may be more important for survival of certain types of cells, although the studies reporting negative results used human and mouse neurons (this work), HEK293 cells (61), Drosophila (63), *S. cerevisiae* (64), and *S. pombe* (51). The yeast studies are informative, as they show that overexpressed human or *S. pombe* DJ-1 does not rescue the greatly increased sensitivity to exogenous MG that results from knockout of Glo1. By contrast, these same studies showed that Hsp31 proteins, which are more active glutathione-independent glyoxalases in the DJ-1 superfamily, can rescue viability of Glo1-deficient yeasts (51,64). Hsp31 has a k_cat_/K_M_ for MG of ~10^3^-10^4^ M^-1^ s^-1^, which is 10-100-fold higher than DJ-1 but still 1000-10000-fold lower than Glo1. Therefore, the ability of Hsp31 to rescue MG sensitivity in ΔGLO1 yeast shows that these complementation experiments are sensitive enough to detect glyoxalase activities that are several orders of magnitude lower than Glo1. In light of our present results and prior independent negative reports, DJ-1’s glyoxalase activity does not appear to be physiologically relevant for overall cell viability when challenged with exogenous MG. Our results do not directly address a potential role for DJ-1 in protecting specific proteins from glycation, which has been reported for histones (57,62). It is speculatively possible that DJ-1’s weak glyoxalase activity may be especially important for sensitive proteins or specific cellular compartments that are not addressed in this global study of cellular glycation.

Very recently, DJ-1 and its close homologs were shown to strongly protect proteins and metabolites against glycerate and phosphoglycerate modifications (37). In that study, DJ-1 was proposed to use Cys106 to open a reactive cyclic 1,3 phosphoglycerate metabolite that may be spontaneously formed from 1,3 bisphosphoglycerate (i.e. DJ-1 is a cyclic 1,3 phosphoglycerate phosphodiesterase). This enzymatic activity is difficult to directly assay owing to the instability of the presumptive substrate so we did not investigate it in this study. However, we note that such an activity may provide a functional explanation for the tendency of the Arg28/Arg48 motif near the DJ-1 active site to bind tetrahedral anions (38). The presence of this anion-binding di-arginine motif is more consistent with DJ-1 acting on a phosphate-containing substrate than with MG being its primary substrate. As we discuss above for DJ-1 glyoxalase activity, mutation of Glu18 could be used to interrogate the importance of a DJ-1 cyclic 1,3 phosphoglycerate phosphodiesterase activity relative to other potential roles of oxidized isoforms of the protein, which we predict would be affected differently by E18N/D/Q mutations. Finally, it is intriguing (though possibly coincidental) that both glyoxalase and cyclic 1,3 phosphoglycerate phosphodiesterase activities act on triose phosphate-derived metabolites resulting from glycolysis, which is restricted to the cytosol in eukaryotic cells.

## Material and Methods

### Protein Expression and Purification

The gene for human DJ-1 was cloned between the NdeI and XhoI sites of *Escherichia coli* expression vector pET15b and the C106S, E18D, and E18N mutants were generated by site-directed mutagenesis and previously described (38). The constructs were transformed into E. coli BL21 (DE3) cells and protein expression and purification were performed as described previously (80). Briefly, the protein was purified by Ni^2+^ metal affinity chromatography, the hexahistitdine tag was removed from recombinant DJ-1 by thrombin cleavage, and the protein was passed through Hi-Q anion exchange column (Bio-Rad Laboratories, Hercules, CA, USA) to remove minor nucleic acid contamination. Thrombin was removed using benazamidine-sepharose 4 resin (Cytiva catalog number 17512310). All proteins ran as a single band on overloaded Coomassie-stained SDS-PAGE and were concentrated using a 10-kDa cutoff centrifugal concentrator (Millipore) to 20 mg/ml in storage buffer (25 mM HEPES pH 7.5, 100 mM KCl, 2-5 mM DTT). DJ-1 concentration was determined by the absorbance at 280 nm using a calculated extinction coefficient at 280 nm of 4400 M^−1^ cm^−1^. The purified protein in storage buffer was snap-frozen in liquid nitrogen and stored at −80 °C.

DJ-1 was oxidized at Cys106 as described previously (80). Briefly, 0.25 mM DJ-1 in storage buffer was dialyzed against PBS (pH 7.4), followed by addition of H_2_O_2_ (Invitrogen) to a final concentration of 1.75 mM (i.e. a 7:1 molar ratio of H_2_O_2_ to DJ-1 monomer) and the mixture was incubated on ice for 45 min. Unreacted H_2_O_2_ was removed by Bio-Gel P-6 desalting resin. Prior work has shown that this procedure selectively oxidizes Cys106 in human DJ-1 (27,70,80). Electrospray mass spectrometry using an Agilent 1200 LC system (Agilent Technologies, Santa Clara, CA, USA) with the electrospray ionization (ESI) source of a Bruker Solarix – 70 hybrid Fourier transform mass spectrometer (Redox Biology Center (RBC) Mass Spectrometry Core Facility, the University of Nebraska-Lincoln) confirmed that DJ-1 was purified in the fully reduced form and that H_2_O_2_ oxidation increased the intact mass of DJ-1 by 32 a.m.u, consistent with Cys106-SO_2_^-^ formation. In addition, preparations of DJ-1 were tested for proper folding by crystallization and X-ray crystal structure analysis. All preparations of the protein crystallize readily and produce crystal structures that are essentially identical (Ca RMSD ~0.05 Å) to prior human DJ-1 structures deposited in the Protein Data Bank (e.g. accession code 1P5F, 5SY6).

### In vitro deglycase and glyoxalase assays

Commercial methylglyoxal (MG) is known to be contaminated with several species that may interfere with kinetic measurements. Therefore, MG was purified from pyruvaldehyde-1-dimethyl acetal as previously reported (67) and stored frozen in small aliquots at −80 °C until needed. 10 mM N-acetyl-cysteine (Alfa Aesar catalog number A15409), glutathione (Acros Organics catalog number 120000250), or coenzyme A (Affymetrix catalog number 13787) were mixed with 10 mM of freshly thawed methyglyoxal and incubated at 25°C until the absorption at 288 nm was stable (~1 hour), indicating that hemithioacetal formation was at equilibrium.

DJ-1 was dialyzed against degased PBS buffer (137 mM NaCl, 2.7 mM KCl, 10 mM Na_2_HPO_4_, 1.8 mM KH_2_PO_4_ pH 7.4) immediately before the kinetic measurements. This buffer was chosen because it was used by both Richarme et al. (55) and Pfaff et al. (63), thereby facilitating direct comparison of our results to prior reports. We note that DJ-1 is reported to be less active in PBS than in other buffers (79), which might be relevant for reconciling the low glyoxalase activity of DJ-1 in vitro with its observed effects in vivo. Andreeva et al. recently reported new equilibrium constants for the formation of MG-hemithioacetals, which are substantially lower than prior values (61). Therefore, not all of the added MG and thiol react to form the hemithioacetal species (79). Initial concentrations of the NAC-MG and GSH-MG hemithioacetal substrates were determined using the updated molar extinction coefficient of 300 M^-1^cm^-1^ and 250 M^-1^cm^-1^, respectively (61). The CoA-MG hemithioacetal molar extinction coefficient has not been reported, so it was presumed to be the same as GSH-MG. The reaction was initiated by the addition of DJ-1 (wild-type, Cys106-SO_2_^-^, C106S, E18N and E18D) to 10 μM final concentration. The A_288_ was measured for six minutes at 25 °C using a Cary50 UV-visible spectrophotometer (Varian, Palo Alto, CA, USA) and always showed a linear decrease over this timeframe. The slope of the best-fit line to the measured decrease in A_288_ signal was used to calculate the initial rate (V0) of hemithioacetal consumption. Because the hemithioacetal spontaneously degrades in aqueous solutions, a buffer-alone baseline was measured in order to correct for non-catalytic loss of 288 nm signal. The difference between the buffer-alone and buffer-enzyme curve was taken to reflect DJ-1 activity. All measurements were repeated at least three times and average values with associated standard deviations are reported.

DJ-1 glyoxalase activity was measured using a coupled assay of L-lactate oxidase and Amplex Red/horseradish peroxidase (HRP). In this assay, DJ-1 glyoxalase activity generates L-lactate (71), which is oxidized by L-Lactate oxidase and molecular oxygen to generate pyruvate and H_2_O_2_. The liberated H_2_O_2_ and Amplex Red are co-substrates for HRP, generating fluorescent resorufin. The rate of H_2_O_2_ generation was monitored using the Amplex Red Hydrogen Peroxide/Peroxidase Assay Kit (Invitrogen). Various concentrations of purified MG were added to the Amplex Red working solution (including Amplex Red reagent and horseradish peroxidase (HRP)), 0.4 units of L-lactate oxidase (Sigma, catalog number L9795), in reaction buffer (50 mM sodium phosphate, pH 7.4, as provided in the kit) at 25°C. The reaction was initiated by adding human DJ-1 to a final concentration 0.1 μM. H_2_O_2_ generation was measured for 5 minutes using a Cary Eclipse spectrofluorimeter (Varian, Palo Alto, CA, USA) with an excitation wavelength at 540 nm and emission wavelength of 590 nm. The measured rates are linear in DJ-1 concentration, confirming that the coupling enzymes were not rate-limiting. All reaction rates were linear over this timeframe and the slope of the best-fit line was converted to H_2_O_2_ concentration using a standard curve generated with known concentrations of H_2_O_2_.

### DJ-1 glyoxalase/deglycase kinetic simulations and data fitting

Data were fit using KinTek global kinetic explorer software (version 10.1.8.beta). The KinTek software simulates reactions by automatically deriving the corresponding system of rate equations, solving them using numerical integration, and fitting kinetic data using non-linear least squares (81). Initial conditions were set using the experimental values. Loss of MG-NAC hemithioacetal was simulated using a molar extinction value at 288 nm of 300 M^-1^cm^-1^, along with the mechanism and rate constants described in Table 2 using experimental conditions described above. The simulation recapitulated the experimental protocol, simulating initial hemoithioacetal formation by mixing 10 mM of MG and NAC for 2 hrs, diluted as indicated in Figure 4, and mixed with DJ-1 (10 μM) for 2 minutes prior to data collection at 288 nm. The equilibrium constant for reaction 2 was fixed using a previously determined value of 500 M^-1^ (61). The rate constants for reaction 1 and 2 were adjusted to globally fit all progress curve data, where best-fit rate constants can be found in Table 2. Confidence intervals for parameters were obtained by the KinTek software (82) by finding rates constants that produce a 0.83 relative error or chi squared ratio 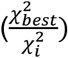 to the best-fit found in Table 2. DJ-1-catalyzed L-lactate formation was monitored through a coupled lactate oxidase and amplex red assay as described above. Lactate oxidation and subsequent formation of H_2_O_2_ to produce fluorescence were made non-rate limiting in the simulation by increasing the rate constant for formation of fluorescent product to an arbitrary value of 3.9 x 10^4^ min^-1^. Simulation of the fluorescent product (Figure 4B) using a linear fluorescent coefficient was based on a H_2_O_2_ Amplex Red standard curve but then optimized to 3.63 x 10^7^ M^-1^ in the fitting.

### Culture of M17 neuroblastoma cell lines

Clonal M17 cell lines stably expressing different control and DJ-1 shRNA sequences were manufactured as described previously (35). Cells were cultured to confluency in Opti-MEM (ThermoFisher, catalog number 31985088) + 10% fetal bovine serum (Gemini Bio-Products, catalog number 900-108) + 5 μg/mL Blasticidin (ThermoFisher, catalog number A1113903). For treatment experiments, control and DJ-1 shRNA M17 cells (n=4) were treated with either regular media or buthionine sulfoximine (43 mM) for 8hr. Following treatment, cells were processed for DNA, RNA, and protein isolation (see below). Treatment concentrations used were the IC_50_ determined by toxicity following 24 hr treatment exposure using 0, 0.1, 1, 10 and 100 mM for buthionine sulfoximine (83).

### Murine primary neuronal cultures

Cortical neuron-enriched primary cultures were prepared from wildtype and DJ-1 knockout mouse pups between postnatal days 1-2 and cultured in Basal medium eagle (+ 1x B-27 minus vitamin A, 1x N2 supplement, 0.5 mM glutamine, 45% glucose, 1x Penicillin-Streptomycin).

### Monolayer forebrain neuronal differentiation from human iPSC

A commercial iPSC line (ThermoFisher, catalog number A18945) and the DJ-1 knockout iPSC line HT-188 (72) were grown on a layer of MEFs (MTI-GlobalStem, catalog number GSC-6201G) in Essential 8TM media (ThermoFisher, catalog number A1517001) until 100% confluent. Confluent cells were transitioned to N3 media (50% DMEM/F12, 50% Neurobasal with 1x Penicillin-Streptomycin, 0.5x B-27 minus vitamin A, 0.5x N2 supplement, 1x Glutamax, 1x NEAA, 0.055 mM 2-mercaptoethanol and 1μg/ml Insulin) plus 1.5 μM Dorsomorphin (Tocris Bioscience) and 10 μM SB431542 (Stemgent) daily for 11 days. On days 12-15, cells were fed each day with N3 without Dorsomorphin and SB431542. On days 16-20, N3 was supplemented with 0.05 μM retinoic acid. On day 20, cells were split 1:2 with trypsin and seeded with ROCK inhibitor onto Poly-L-ornithine (Sigma), fibronectin and laminin coated plates. From days 21 to time of extraction, cells were fed with N4 media (same as N3 plus 0.05 μM Retinoic acid, 2 ng/ml BDNF and 2 ng/ml GDNF).

### DJ-1 knockout mice

Male and female WT (n=5) and DJ-1 knockout (n=6) mice (C57BL/6) were given access to food and water ad libitum and housed in a facility with 12-h light/dark cycles. Mice were sacrificed between 20-22 months of age and brains were flash frozen and stored at −80°C. Brains were homogenized into a frozen powder using liquid nitrogen and separated for DNA, RNA, and protein extractions.

### RNA extraction for glycation analysis

All cell lines were seeded at ~1×10^6^ per 12 well with n=4 wells used for RNA, DNA, or protein extraction. Cells were lysed using 0.4 mL TRIzol™ reagent (ThermoFisher, catalog number 15596026) and 0.08 mL chloroform. Following centrifugation for 15 minutes at 4°C at 12,000xg, the top aqueous layer was transferred to a new Eppendorf tube. RNA was precipitated with the addition of 0.2 mL isopropanol followed by centrifugation for 10 minutes. RNA pellet was washed with 0.4 mL 75% ethanol and centrifuged for 5 minutes at 7,500xg. Supernatant was discarded and pellet was air dried then resuspended in 20 μL RNase-free water. To complete solubilizing the RNA, incubation in water bath at 55°C occurred for 15 minutes. RNA from brain samples were processed using the same protocol. RNA concentration was quantified using spectrophotometer.

### DNA extraction for glycation analysis

Cells were rinsed, scraped, and pelleted using sterile PBS. Cells were resuspended in equal volumes PBS and phenol:chloroform:isoamyl (25:24:1) and spun for 5 minutes at 16,000xg. Top aqueous layer was transferred to a new Eppendorf tube. DNA was precipitated by adding 0.5x volume of 7.5 M ammonium acetate and 3x volume of 100% ethanol then stored at −20°C overnight. DNA was pelleted by centrifugation for 30 minutes at 4°C at 16,000xg. Supernatant was removed and pellet was washed with 70% ethanol. Following centrifugation, supernatant was removed, pellet was air dried and resuspended in 100 μL autoclaved water. DNA from brain samples was isolated using the DNeasy Blood and Tissue kit (Qiagen, 69504). DNA concentration was quantified using spectrophotometer.

### Protein extraction for glycation analysis

Cells were scraped and lysed with lysis buffer (1x cell lysis buffer (Cell Signaling Technology, 9803), 1x protease inhibitor (Sigma, 4693159001), and 1x phosphatase inhibitor (ThermoFisher, 78427)). Lysed cells were set to rotate for 30 minutes at 4°C then spun at >20,000xg for 8 minutes at 4°C. Supernatant was transferred to a new Eppendorf tube and protein concentration measured using the Pierce™ 660 nm protein assay. Protein from brain samples were extracted using a similar protocol.

### HPLC analysis of MG reaction with dG in the presence of DJ-1

MG was synthesized as previously described (84) and reacted (5 mM) with dG (0.5 mM) in 50 mM sodium phosphate, pH 7.5 for 1 hr at 37 °C. DJ-1 (5 μM) was added for an additional 24 hr. Reactions were also performed in which DJ-1 was added concomitantly with MG and dG. Following incubation, reactions were analyzed on an Agilent 1100 HPLC as previous described (67). Reactions with dinitrophenylhydrazine (DNPH) were performed by making a 10 mM DNPH stock in 2.5 M HCl with dilution to 5 mM in H_2_O. DNPH (5 mM) was added to reactions for 15 minutes followed by addition of 2.5 M NaOH prior to injection on HPLC. Absorbance from 210-510 nm was monitored.

### Mass Spectrometric analysis of CEdG, CEG, and CEL from WT and DJ-1 KD cells

Standards were synthesized as previously described (67). CEL-d4 standard was obtained from Santa Cruz Biotechnology (sc219424). CEL was obtained from Abcam (ab145095). DNA and RNA was isolated as described above, spiked with 5 ng/mL ^15^N_5_-CEdG or ^15^N_5_-CEG and digested as previously described (84). Protein was isolated as described above and up to 50 μg of protein per sample was spiked with 5 ng/mL of CEL-d4 and Lys-d4. Samples was incubated in 6 M HCl for 18 hr at 110 °C and then dried under nitrogen with incubation at 80 °C, resuspended in 50 μL 0.5% HFBA, and then analyzed using LC-MS/MS. CEdG and CEG were analyzed by injection onto an Agilent Zorbax SB-C18 column (2.1×50mm, 1.8 μm) with mobile phases A: H_2_O + 0.1% formic acid and B: acetonitrile + 0.1% formic acid at 0.4 mL/min. The following gradient was used to separate analytes: 3%-10% B from 0-4.5 min, 10%-97% B from 4.5-4.8 min, 97% B from 4.8 to 5.2 min, 97% to 3% B from 5.2 to 5.7 min. CEL was analyzed on this same column with an isocratic flow of 10% ACN + 0.1% formic acid at 0.2 mL/min. The following mass transitions were monitored: CEdG *m/z* 340 to 224, ^15^N_5_-CEdG *m/z* 345 to 229, CEG *m/z* 356 to 224, and ^15^N_5_-CEG *m/z* 361 to 229. CEL *m/z* 219 to 130 CEL-d4 *m/z* 223-134, Lys *m/z* 147-84, and Lys-d4 *m/z* 151 to 88. Analytes were quantified by fitting to a standard curve and the normalization to dG or guanosine (measured as described below), or lysine, as appropriate.

### HPLC quantitation of dG and Guanosine

Following mass spectrometric quantitation of CEdG and CEG, dG and guanosine were quantified on an Agilent 1100 HPLC using an Atlantis T3 column 4.6 x 150 mm, 5 μm. The mobile phases were A: H_2_O + 0.1% formic acid and B: ACN + 0.1% formic acid with a flow rate of 0.426 mL/min. The following gradient was used: 0-9% B, 0-20 min; 9-9.5% B, 20-55 min; 9.5%-90% B, 55-60 min and hold 60-70 min. 90%-0% B, 70-75 min; 0% B, 75-80 min. The area under the curve was manually integrated and then fit to a standard curve to determine dG or G concentration.

### Statistical analysis

Comparisons between various groups were performed using two-way ANOVA with Tukey’s multiple comparisons test and an alpha value of 0.05 as implemented in Prism 9.3.0 (GraphPad software).

## Acknowledgements/Conflict of Interest Disclosure

We thank Dr. Javier Seravalli (Redox Biology Center Mass Spectrometry Core Facility, University of Nebraska) for assistance with mass spectrometry of recombinant DJ-1. This research was supported in part by the Intramural research Program of the National Institutes of Health (NIH; Bethesda, MD, USA), National institute on Aging. MAW acknowledges support from NIH R01GM139978, JT R01CA176611. Work performed in the Mass Spectrometry Shared Resource at City of Hope is supported by the National Cancer Institute of the National Institute of Health under award number P30 CA033572. The authors declare no conflicts of interest that would affect the objectivity of this study.

