## Supplemental material for "DJ-1 glyoxalase activity makes a modest contribution to cellular defense against methylglyoxal damage in neurons"

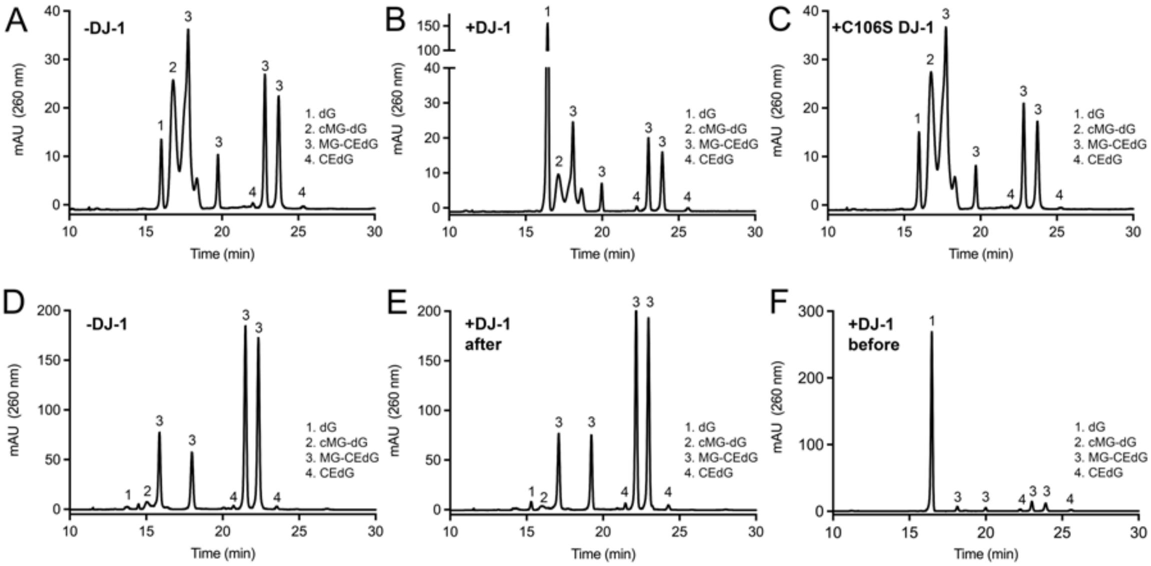


**Supplemental Figure S1**. **HPLC analysis of the effect of DJ-1 on dG glycation in vitro.** Raw HPLC elution profiles with peaks labeled by species are shown for all conditions in Fig 2A,B. In (E), "+DJ-1 after" is the result of adding DJ-1 after preincubation of MG and dG. In (F), "+DJ-1 before" is the result of adding DJ-1 at the same time and MG and dG.

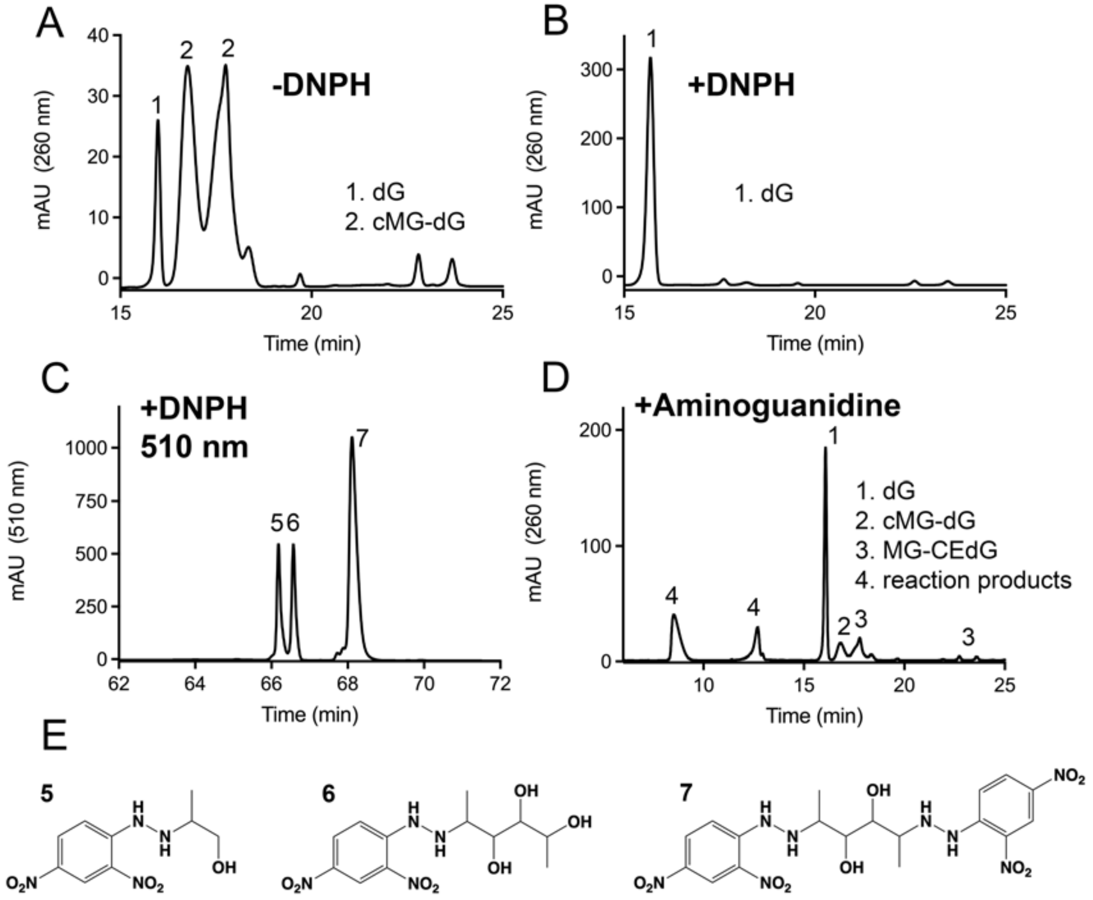


**Supplemental Figure S2. Aldehyde scavengers reduce glycation products similarly to DJ-1.** (A) HPLC elution profile of dG glycation by MG in the absence of DNPH. (B) Addition of DNPH with MG results in predominantly unmodified dG, similar to the effect of DJ-1 in Fig. 2A,B and Supplemental Fig. S1F. (C) Incubation of DNPH with MG creates several species that absorb at 510 nm. (D) Aminoguanidine has similar effects to DNPH (B) and DJ-1 (Fig. 2A,B and Supplemental Fig. S1F) on preventing dG glycation. (E) Chemical structures for the 510 nm-absorbing species formed by DNPH and MG, with numbers corresponding to the peaks in (C).

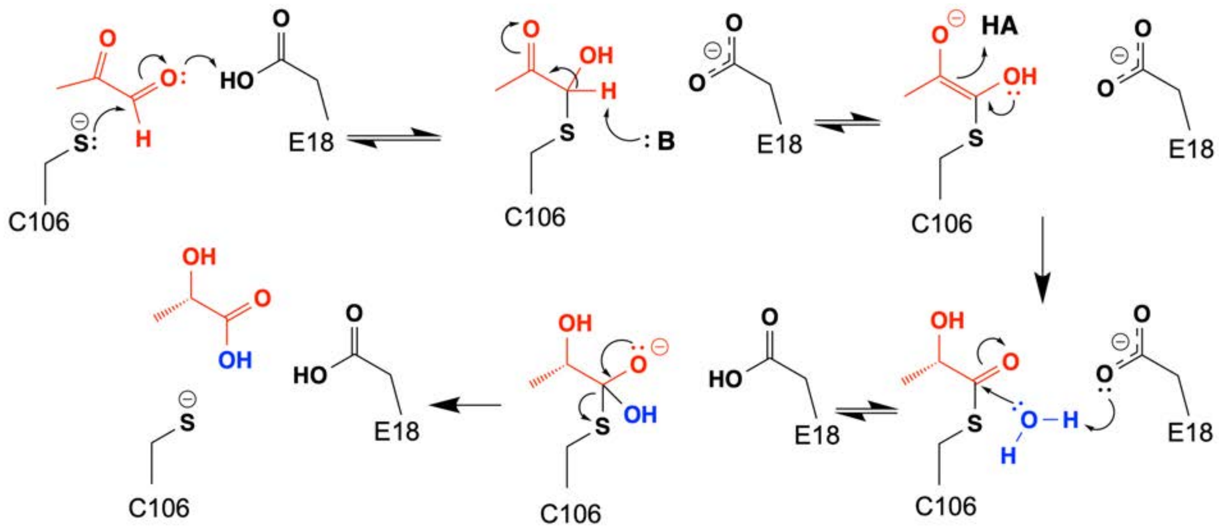


**Supplemental Figure S3. Proposed mechanism for DJ-1 glyoxalase activity.** Electron flow is shown with curved arrows, with MG in red and water in blue. Direction of the reaction is shown with straight arrows, with some steps presumed reversible shown in double arrows. We note that the identity of the general base B and the general acid HA are not known but may represent a protonation/deprotonation cycle of Glu18, although this is speculative.

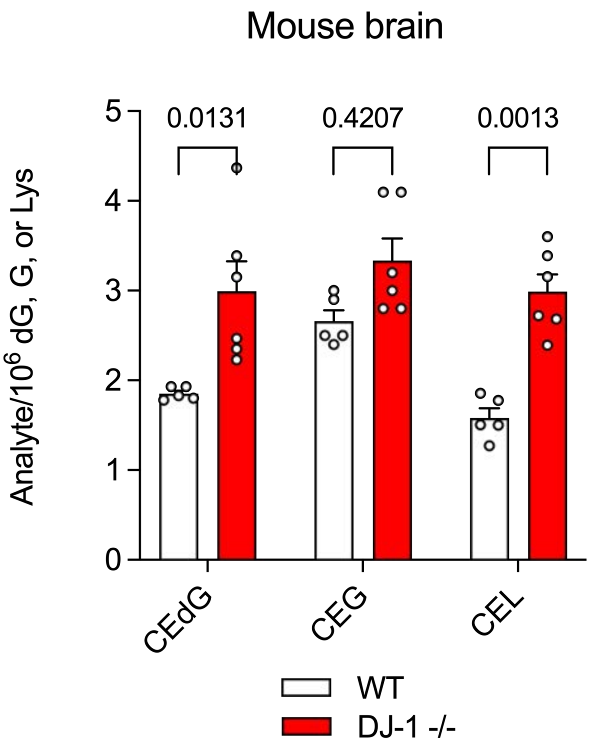


**Supplemental Figure S4. DJ-1 slightly decreases cellular concentrations of irreversible glycation products in whole mouse brain**. In all panels, isotope-dilution mass spectrometry was used to obtain relative concentrations of modified vs. unmodified dG, G, or Lys. Two-way ANOVA was used for statistical analysis with p-values shown. Small but consistent elevations in glycated products were observed in whole brains from DJ-1$^{-/-}$ mice compared to WT controls. Each measurement is shown as a circle with standard error of the mean shown in error bars.
